# Predicting Drug Synergy and Antagonism from Genetic Interaction Neighborhoods

**DOI:** 10.1101/050567

**Authors:** Jonathan H. Young, Edward M. Marcotte

**Affiliations:** Institute for Computational Engineering and Sciences, Center for Systems and Synthetic Biology, The University of Texas at Austin, Austin, Texas, USA; Center for Systems and Synthetic Biology, Institute for Cellular and Molecular Biology, Department of Molecular Biosciences, The University of Texas at Austin, Austin, Texas, USA

**Keywords:** Drug interactions, Synergy, Antagonism, Yeast

## Abstract

Although drug combinations have proven efficacious in a variety of diseases, the design of such regimens often involves extensive experimental screening due to the myriad choice of drugs and doses. To address these challenges, we utilize the budding yeast Saccharomyces cerevisiae as a model organism to evaluate whether drug synergy or antagonism is mediated through genetic interactions between their target genes. Specifically, we hypothesize that if the inhibition targets of one chemical compound are in close proximity to those of a second compound in a genetic interaction network, then the compound pair will exhibit synergy or antagonism. Graph metrics are employed to make precise the notion of proximity in a network. Knowledge of genetic interactions and small-molecule targets are compiled through literature sources and curated databases, with predictions validated according to experimentally determined gold standards. Finally, we test whether genetic interactions propagate through networks according to a “guilt-by-association” framework. Our results suggest that close proximity between the target genes of one drug and those of another drug does not strongly predict synergy or antagonism. In addition, we find that the extent to which the growth of a double gene mutant deviates from expectation is moderately anti-correlated with their distance in a genetic interaction network.

## Introduction

Drug combinations have an established history in treating disease, dating to the MOPP regimen for Hodgkin’s lymphoma in the 1960s to highly active antiretroviral therapy (HAART) for HIV in the 1990s [Hammer et al., 1997, DeVita and Chu, 2008]. In combating antibiotic resistance, combination regimens have proven effective and are actively under continued development [Worthington and Melander, 2013]. Yet in designing combination therapies, it is not immediately clear which drugs and doses to group together; there are simply a myriad of possible choices and the combinatorial space quickly grows unwieldy. As a result, any computational technique to either guide readily testable candidates or reliably predict the effect of drug combinations would be desirable. In this study, using the budding yeast *Saccharomyces cerevisiae* as a test platform, we determine whether the effect of drug pairs can be predicted from genetic interactions between their target genes.

The effect of a drug combination can be classified as synergistic, antagonistic, or additive. Two drugs are synergistic if they cause a significantly greater growth defect than expected, based on the effect of each drug individually. Antagonism is similar, although the effect is far more pronounced growth than expected. Drug additivity implies that no interaction exists between the agents, and the resulting phenotype is the sum of each drug’s individual effect. There is more than one choice of a null model that defines the “expected effect” - commonly used models include Loewe additivity and Bliss independence [Loewe, 1953, Bliss, 1939, Yeh et al., 2009].

Previous studies to uncover genetic interactions as a mechanism underlying drug combinations have involved exhaustive screening of a number of small-molecule chemical compounds. An examination of 200 compound pairs administered in *Saccharomyces cerevisiae* found 38 of them to be synergistic, but genetic interactions were determined to be responsible for only 14 of those 38 [Cokol et al., 2011]. Another study screened all possible pairs of 128 compounds from a chemically diverse library to experimentally deduce synergy and antagonism, thereby establishing a validation set. Moreover, a model based directly on chemical-genetic and genetic interactions had low predictive power for synergy or antagonism, but combining naive Bayes and random forests trained on additional features led to successful predictions [Wildenhain et al., 2015].

We hypothesized that the proximity in a genetic interaction network between one drug’s target genes and another drug’s targets controls the degree to which the drug pair is synergistic or antagonistic. In particular, rather than considering only direct interactions between genes, our approach factored in whether a gene is within a neighborhood of (though not necessarily adjacent to) some other gene in the network. We leveraged knowledge of known small-molecule inhibition targets in *S. cerevisiae* from the Search Tool for Interactions of Chemicals (STITCH) database [Kuhn et al., 2013] and experimentally determined negative and positive genetic interactions [Costanzo et al., 2010]. Finally, predictions of synergy or antagonism were validated against gold standards assembled from the literature.

## Methods

### Negative and positive genetic interaction network

Negative and positive genetic interactions were compiled from a high-throughput yeast synthetic genetic array (SGA) screening dataset [Costanzo et al., 2010]. The intermediate cutoff for the genetic interaction score *∊* was chosen as the threshold for interacting versus non-interacting gene pairs. For the purposes of data processing, the suffixes “_tsq” and “_damp” were removed from gene symbols. Both unweighted and weighted versions of each of the negative and positive genetic interaction networks were assembled. Nodes in the networks correspond to genes and two genes are connected by an edge if they interact according to the intermediate cutoff *∊*. Because a larger magnitude of *∊* indicated stronger genetic interaction, in the weighted networks the edge weights were assigned by reversing the *∊* values. For instance, the strongest genetically interacting pair was assigned an edge weight with the smallest *∊* instead. All edge weights were set to be non-negative.

### Chemical compound targets and gold standards for synergy and antagonism

Two literature sources were used as the gold standard to validate chemical synergy and antagonism predictions [Cokol et al., 2011, Wildenhain et al., 2015]. The inhibition targets in *S. cerevisiae* of chemical compounds identified by CID were assembled from STITCH version 4 [Kuhn et al., 2013]. Only chemical names were available from the Cokol et al. dataset; these were converted to CIDs with PubChemPy https://pypi.python.org/pypi/PubChemPy. SMILES strings from the Wildenhain et al. dataset were also converted to CIDs. Prediction performance was assessed with receiver operating characteristic (ROC) analysis as implemented in scikit-learn [Pedregosa et al.,2011].

### Distances in networks

Distances between all pairs of nodes in unweighted and weighted versions of both the negative and positive genetic network were computed using Dijkstra’s algorithm as implemented in NetworkX [Schult and Swart, 2008]. The distance between two sets *A* and *B* of nodes in an unweighted network was calculated using the earth mover’s metric (EMD) [Rubner et al, 2000]. Here in the 1-dimensional special case, the EMD reduces to differences between cumulative distribution functions [Cohen, 1999]. For the purpose of measuring the distance between two sets of nodes, EMD(*A*, *B*) = Σ_*i∊*No_ **|** *F*_*X*_ref__(*i*) – *F_X_*(*i*) |, where *F*_*X*_ref__ and *F_X_* are the cumulative distribution functions (CDFs) of a reference distribution *X*_ref_ and a random variable *X*. The reference distribution is intended to represent the scenario where every node in *A* is adjacent to some node of *B*. In an unweighted network, the reference probability mass function (pmf) of *F*_*X*_ref__ is defined as

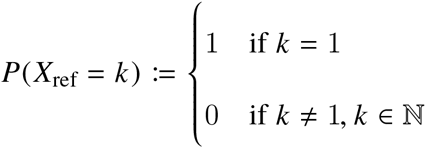

and the pmf for *X* is constructed from the frequencies of all possible node pair distances between *A* and *B* as found from Dijkstra’s algorithm.

In a weighted network, we have

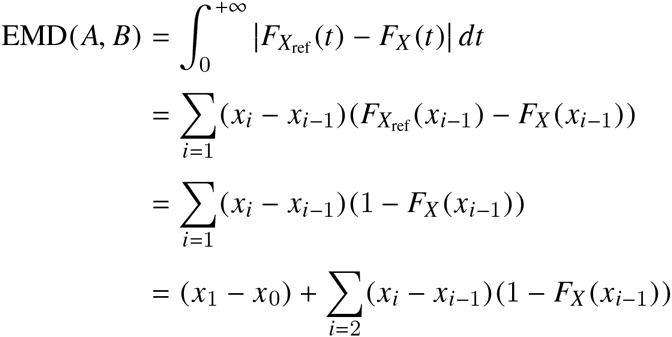

where by choosing *x*_0_ to be the minimum edge weight *P*(*X*_ref_ = *x*_0_) = 1 and 0 elsewhere, and *P*(*X*) is non-zero only for the node pair distances *x*_1_, *x*_2_,… with *x*_0_ ≤ *x*_1_ < *x*_2_ < ···.

### Software availability

Computational analyses were performed with Python version 3.4; scripts and Jupyter notebooks are available under the BSD license at https://bitbucket.org/youngjh/yeast_chem_synergy. All plots were created with Matplotlib and Seaborn [Hunter et al., 2007].

## Results

### Close proximity between drug target genes in the genetic interaction network does not strongly predict synergy or antagonism

We hypothesized that if two chemical compounds are synergistic, then the inhibition target genes of one compound would be close to those of the second compound in a negative genetic interaction network. Similarly, antagonistic compound pairs would have their respective targets near one another in a positive genetic network. Proximity between target genes were assessed in both unweighted and weighted genetic interaction networks. An experimental screen in *S. cerevisiae* provided the gold standard benchmark for testing the synergy hypothesis [Wildenhain et al., 2015]. In this dataset, all possible pairs of 128 chemical compounds were screened, but only 7 compounds had inhibition target genes found in both the negative genetic network and the Search Tool for Interactions of Chemicals (STITCH) database. Thus, there were 21 possible pairs available for validation, three of which exhibited synergy from the screening results. None of the antagonistic compounds in this dataset contained targets listed in STITCH. As shown in Table 1, close proximity of the target genes were only weakly predictive of synergy, according to the area under the curve (AUC) from the receiver operating characteristic (ROC) analysis. The AUC from the unweighted network was reasonably consistent with that from the weighted network.

**Table 1.** Chemical compound pairs were scored and ranked for synergy or antagonism by the distance between their inhibition targets in a genetic interaction network. The predictions were validated through receiver operating characteristic (ROC) analysis with true interactions labeled according to gold standards for synergy and antagonism. In the synergy case, target gene proximity is only marginally more predictive than random for chemical synergy or antagonism.

|  | Synergy | Antagonism |
| --- | --- | --- |
| Unweighted network | 0.61 | 0.41 |
| Weighted network | 0.57 | 0.19 |

Synergy and antagonism prediction performance assessed by AUC, the area under the receiver operating characteristic (ROC) curve.

For the antagonism case, the gold standard was constructed from another experimental screen [Cokol et al., 2011]. Of the 200 pairs screened from 33 compounds, only 10 pairs had compounds whose inhibition targets were both listed in STITCH and in the positive genetic network. Of these 10, 8 were experimentally determined to show antagonism. None of the synergistic compounds in this dataset contained targets listed in STITCH. No evidence was found to suggest that close proximity of target genes was predictive of antagonism (Table 1). In fact, the results suggest that the farther apart one set of target inhibition genes is from those of a second compound in the positive genetic network, the more likely the compound pairs are to be antagonistic. Strikingly, in contrast to the synergy case above, the AUC value from the unweighted network was quite far apart from that of the weighted network.

### Genetic interaction strength is moderately correlated with network distance

One assumption underlying our hypothesis was that any two genes that were not adjacent in the genetic interaction network but within a sufficiently small neighborhood of one another would still express some degree of interaction. Conversely, if the genes were located very far apart, they would essentially not interact at all. To examine the validity of this assumption, we sought to determine the correlation, if any, between a gene pair’s distance in the network and its corresponding strength of genetic interaction. The interaction strength was simply the magnitude of the genetic interaction score |*∊*| from the raw results of the synthetic genetic array (SGA) screening [Costanzo et al., 2010]. The network distance of a gene pair was once again the distance computed from Dijkstra’s algorithm as described above, such that smaller distances implied stronger interaction and consequently larger |*∊*|. Therefore, we expected to observe negative correlations for both negative and positive genetic. As shown in Figure 1, indeed the Spearman’s rank correlation correlation is in fact moderately negative and statistically significant.

**Figure 1.**
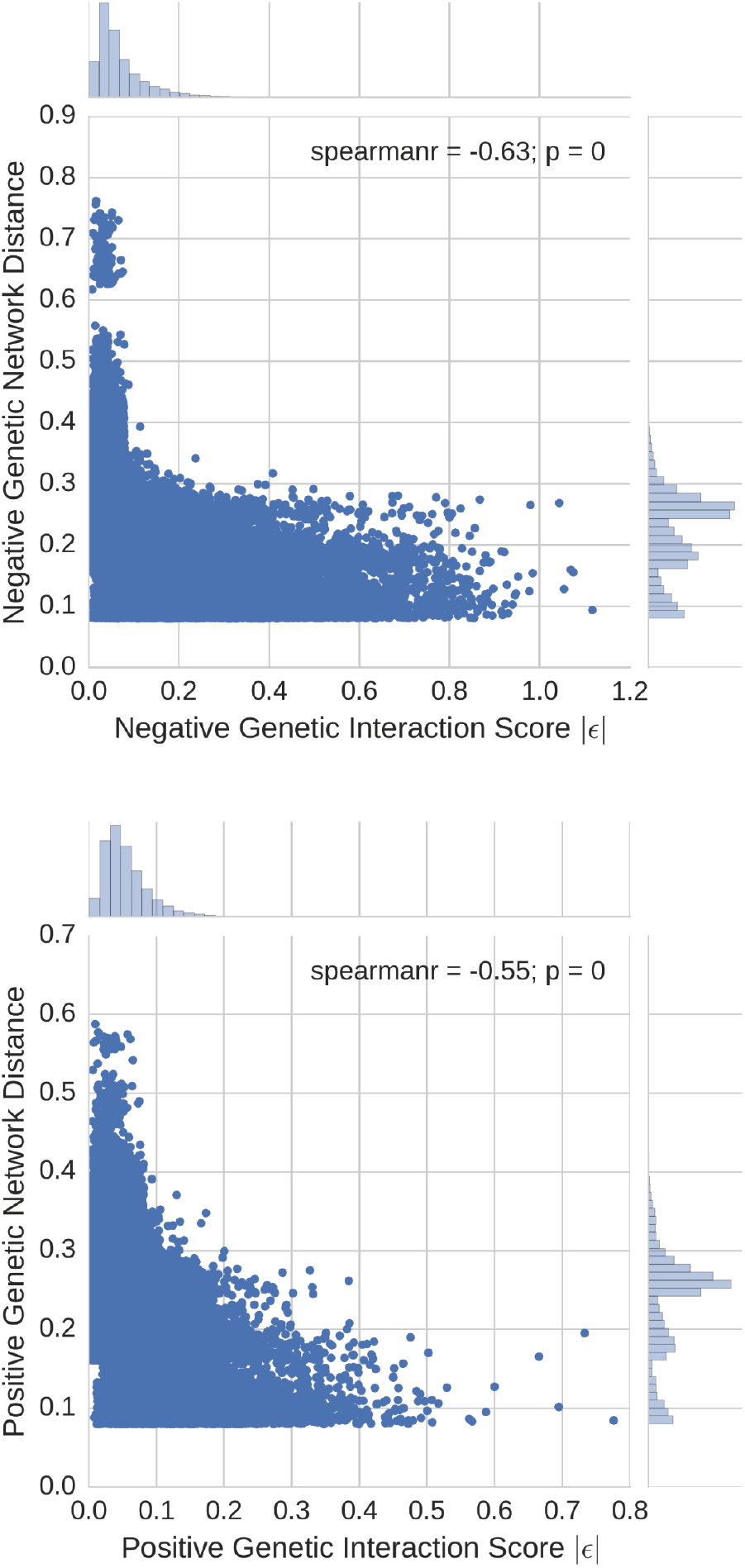
The magnitude of the genetic interaction score *∊* is moderately anti-correlated with gene network distance. Thus, the greater the growth deviation from expectation of a double mutant, the closer the two genes are in the genetic interaction network.

## Discussion

Our results suggest that there is no evidence to support the claim that synergy or antagonism arises when the target genes of one chemical compound are close in a genetic interaction network to those of another compound. We confirmed previous results that such drug interactions are not directly mediated through genetic interactions [Cokol et al., 2011, Wildenhain et al., 2015], and also showed that neighborhoods of genetic interactions are neither a contributing factor as well. In the process, we presented an application of distance measures satisfying the mathematical definition of a metric to quantify proximity between sets of nodes in gene networks. Prediction performance was measured through AUC due to its robustness to unbalanced data in positive versus negative class labels [Jeni et al., 2013].

It is particularly notable that the gold standard for synergy produced results different than those from the antagonism gold standard. One potential contributing factor is that the benchmark derived from Cokol et al. used the Loewe additivity model [Loewe, 1953, Tallarida, 2006] to determine synergy and antagonism, while Wildenhain et al. instead utilized Bliss independence [Bliss, 1939]. The Bliss theory is closer to the multiplicative fitness model employed in calling negative and positive genetic interactions, which was defined as *∊_ij_* = *f_ij_* – *f_i_ f_j_* with *f_ij_* equal to the double mutant fitness and *f_i_*, *f_j_* as the single mutant fitness scores [Costanzo et al., 2010].

The moderate correlation between genetic interaction strength and network distance goes some way towards supporting the results from the synergy gold standard, where AUCs of 0.57 and 0.61 were attained. In any case, the weak correlation implies that genetic interactions cannot be reliably identified through “guilt-by-assocation” in the network. It should be noted that both datasets used to benchmark prediction performance were highly imbalanced, thus reflecting the need for even more data on which chemical compounds are synergistic or antagonistic, and which genes are inhibited by the compounds of interest. Yet despite the relatively limited data available to construct gold standards, our results and those of others indicate that a more nuanced mechanism beyond genetic interactions of target genes is responsible for explaining effects of chemical compound interactions.

